# Local sleep in songbirds: Different simultaneous sleep states across the avian pallium

**DOI:** 10.1101/2023.10.20.563239

**Authors:** Hamed Yeganegi, Janie M. Ondracek

**Author notes:** Correspondence: Janie M. Ondracek, Technical University of Munich, TUM School of Life Sciences, Chair of Zoology, Liesel-Beckmann-Str. 4, 85354 Freising-Weihenstephan, Germany.

## Abstract

Wakefulness and sleep have often been treated as distinct and global brain states. However, an emerging body of evidence on the local regulation of sleep stages challenges this conventional view. Apart from unihemispheric sleep, the current data that support local variations of neural oscillations during sleep are focused on the homeostatic regulation of local sleep, i.e., the role preceding awake activity. Here, to examine local differences in brain activity during natural sleep, we recorded the electroencephalogram (EEG) and the local field potential (LFP) across multiple sites within the avian pallium of zebra finches without perturbing the previous awake state. We scored the sleep stages independently in each pallial site and found that the sleep stages are not pallium-wide phenomena but rather deviate widely across electrode sites. Importantly, deeper electrode sites had a dominant role in defining the temporal aspects of sleep state congruence. Altogether, these findings show that local regulation of sleep oscillations also occurs in the avian brain without prior awake recruitment of specific pallial circuits and in the absence of mammalian cortical neural architecture.

## Introduction

Sleep is a phenomenon that has been documented across animal species (Cirelli & Tononi, 2008; Siegel, 2008) and can be defined on at least two levels: i) the behavior of the whole organism and ii) the patterns of neural activity in the brain (Vyazovskiy & Harris, 2013). Behavioral sleep is often simply defined as a state of quiescence and reduced responsiveness to the environment. In contrast, electrophysiological sleep is characterized by the complex spatiotemporal patterns of neural activity that occur during behavioral sleep.

In mammals, the sleeping brain generally oscillates between two principle states: rapid eye movement (REM) sleep and non-REM (NREM) sleep. These two states of sleep have also been documented in numerous non-mammalian animals, including birds (Canavan & Margoliash, 2020; Libourel et al., 2023; Low et al., 2008; Yeganegi & Ondracek, 2023), reptiles (Albeck et al., 2022; Libourel et al., 2018; Shein-Idelson et al., 2016), zebra fish (Leung et al., 2019), and cephalopods (Pophale et al., 2023). Similar to mammals, REM sleep-like states in these animals are characterized by low amplitude and aperiodic electrical activity, whereas NREM-like sleep is characterized by the emergence of periodic “slow” wave activity (1-4 Hz oscillations) during slow wave sleep (SWS).

REM sleep and NREM sleep states are conventionally described as unitary “global” phenomena, affecting the whole brain uniformly and simultaneously. However, recent work on “local” sleep in both humans (Bernardi et al., 2015; Huber et al., 2004; Nir et al., 2011) and rodents (Funk et al., 2016; Rector et al., 2005; Vyazovskiy et al., 2011) has challenged this understanding, suggesting that brain activity is much more complex and compartmentalized.

In the field of sleep research, the phrase “local sleep” has been used to describe a variety of different phenomena. In mammals, the term “local sleep” most commonly refers to two distinct concepts.

(1) Local sleep may refer to the use-dependent, or homeostatically influenced, differences in slow wave activity (SWA) observed during NREM sleep (Hanlon et al., 2009; Huber et al., 2004; Kattler et al., 1994; Lesku et al., 2011; Vyazovskiy & Tobler, 2008). Here “local” refers to the observation that after awake unilateral experience (e.g. motor learning, sensory stimulation), a “local” SWA asymmetry results across the hemispheres during subsequent sleep phases, with the hemisphere or brain area contralateral to the stimulation displaying larger SWA values. This phenomenon is thought to result from the unilateral and activity-dependent upregulation of genes involved in long-term potentiation resulting in a “local” increase in sleep requirement (Hanlon et al., 2009).

(2) Local sleep may describe other disparities of sleep states, ranging from i) the intrusion of sleep-like activity in the cortex of awake and behaving animals (Vyazovskiy et al., 2011), where “local” sleep-like activity appears in one cortical area during awake periods, but not another; ii) the presence of “local” slow waves during REM sleep in mice (Funk et al., 2016); iii) “local” slow waves that are out of phase across brain regions in humans (Nir et al., 2011) and rats (Rector et al., 2005) or iv) the simultaneous co-existence of REM sleep and NREM sleep in different brain structures, such as the hippocampus and neocortex of rats (Durán et al., 2018; Emrick et al., 2016).

It is this second understanding of local sleep that we deal with in this article, and specifically, the exploration of disparate sleep states in smaller, “local” neural assemblies (Krueger et al., 2008), where sleep phenotypes may emerge from the integration and synchronization of local network states (Krueger et al., 2013).

The avian brain provides an important test ground for investigating these local differences during natural sleep. Although most of the theoretical models that examine the generation of neural oscillations depend on cytoarchitecturally regular structures, such as the layered structure of the mammalian cortex (Buzsáki et al., 2012; da Silva, 2010), the avian brain is capable of generating sleep oscillations comparable to that of mammalian slow waves (Canavan & Margoliash, 2020; Low et al., 2008), despite the absence of a layered cortex. By investigating local sleep in the avian brain, we extend the existing knowledge about local sleep across animal species and neural architectures.

In this study, we examined differences in sleep states across two parts of the avian pallium (telencephalon). We recorded 1) superficial EEG signals from the avian hyperpallium, which is homologous to the dorsal pallium (neocortex) of mammals and 2) deeper LFP signals from the avian dorsal ventricular ridge (DVR), which is a pallial structure unique to birds and reptiles. In birds, the DVR is further subdivided into the mesopallium, nidopallium, and arcopallium (Briscoe & Ragsdale, 2018). Importantly, sites in the hyperpallium have been shown to exhibit cortical-like, use-dependent regulation during sleep (Lesku et al., 2011), whereas hippocampal-like sharp-wave ripple activity has been recorded in the DVR of both birds (Yeganegi et al., 2019) and reptiles (Norimoto et al., 2020; Shein-Idelson et al., 2016).

## Methods

### Experimental animals

All experimental procedures were conducted in accordance with the principles of laboratory animal care and were carried out in compliance with the current versions of the German and European laws on Animal Experimentation (Approval # ROB-55.2-2532.Vet_02-18-108: J.M.O.). Additionally, the experiments followed the guidelines set forth by ARRIVE (Animal Research: Reporting of In Vivo Experiments). Ethical approval for the study was obtained from the ethics committee of the Government of Upper Bavaria (Regierung von Oberbayern), which operates in accordance with § 14.1 TierSchG (German animal welfare law). Furthermore, the housing and breeding arrangements for the animals received the approval of the Veterinary Office of Freising, Germany (Approval # 32-568).

Electrophysiological recordings were made in three male zebra finches (*Taeniopygia guttata*), aged 83-91 days post-hatch (dph), all of which exhibited physical signs of maturity as indicated by feather and beak coloring. Details concerning the age and corresponding symbol used for each individual are presented in Table 1. The data analyzed here represent 36 hours of data recorded in 3, 12-hour long sleep recordings sessions from 3 animals.

**Table. 1.** Description of experimental birds used in experiments. Bird ID, age, the corresponding symbol used in the figures, and the number of recording electrodes available are displayed. dph: days post-hatch, RH: right hemisphere. *: data from the left posterior EEG was not used in this bird.

| Bird ID | Age (dph) | Symbol used in this text | # Good EEG electrodes | Medial LFP location |
| --- | --- | --- | --- | --- |
| w027 | 83 | ○ | 4/4 | 1.86 mm (RH) |
| w042 | 90 | + | 3/4 * | 1.74 mm (RH) |
| w044 | 91 | ◇ | 4/4 | 1.25 mm (RH) |

Prior to the experiments, the birds were housed in aviaries with a 12:12-hour light:dark cycle. They were provided with access to a seed mix, millet, sepia bone, and water *ad libitum*. Additionally, fresh vegetables and cooked eggs were given to the birds once a week. The recording chambers were equipped with LED lights, UV bird lights (Bird Systems, Germany), infrared LED panels (850 nm waves), and ventilation to ensure constant air circulation.

### Electrode preparation and positioning

For each animal, an electrode implant was constructed that consisted of 4 EEG electrodes, 4-8 LFP electrodes, a ground and a reference electrode. The preparation and positioning of the EEG electrodes is described elsewhere (Yeganegi & Ondracek, 2023); briefly, EEG electrodes were constructed from silver wire (254 μm bare wire diameter, Science Products GmbH) by melting the tip into a smooth ball of approximately 0.50 mm in diameter. The LFP electrodes were made from 5 mm lengths of coated steel wire with a bare diameter of 127 µm (A-M Systems, Sequim, WA) and pinned to an omnetics connector with small gold EIB pins (Neuralynx Inc, Bozeman, MT, USA). Because the number of LFP electrodes was variable across animals, we chose one LFP electrode to use as the designated LFP electrode in the congruence analysis. This LFP electrode was the most medial LFP electrode, and across animals, it was positioned on average at 1.62 ± 0.32 mm (mean ± std; Supplementary Fig. S1).

### Anesthesia and Surgery

The birds were anesthetized using oxygenated isoflurane gas (EverFlo OPI, Phillips, Netherlands) with a 3-4% concentration during induction and 0.8-1.5% during maintenance. A small tube was used to deliver the anesthetic gas through the bird’s beak via a vaporizer (Isotec 4, Groppler, Germany). To prevent the accumulation of excess anesthetic gas, a scavenger (Scavenger LAS, Groppler, Germany) removed waste anesthetic gas. The bird was positioned on a heating pad with a closed-loop temperature monitoring device (Harvard Apparatus, MA, USA). Throughout anesthesia, body temperature was continuously monitored using an infrared thermometer and remained above 39.5°C.

The anesthetized bird was positioned in a stereotaxic frame (Kopf Instruments, CA, USA) and the head was immobilized. Anesthetic depth assessment was carried out by applying slight pressures on a toe using tweezers. When the toe reflex ceased, a local anesthetic was applied to the scalp (Aspen, xylocaine pump spray), and the feathers on the head were removed with forceps. An incision was made in the scalp along the sagittal midline, and a dental drill (Volvere i7, NSK Europe GmbH, Germany) was used to make small holes in the skull for the (supradural) EEG electrodes (see the schematic in Fig. 1a) and a small craniotomy (1×1.5 mm) for the LFP electrodes. The dura matter was resected, the LFP electrodes were lowered altogether 1.8-2.2 mm deep into the pallium, targeting the mesopallium of the DVR (see the histology sample in Fig. 1B) and secured in place with dental cement (Paladur, Henry Schein Dental, Germany). The EEG electrodes were then positioned over the hyperpallium and secured with dental cement.

**Fig. 1.**
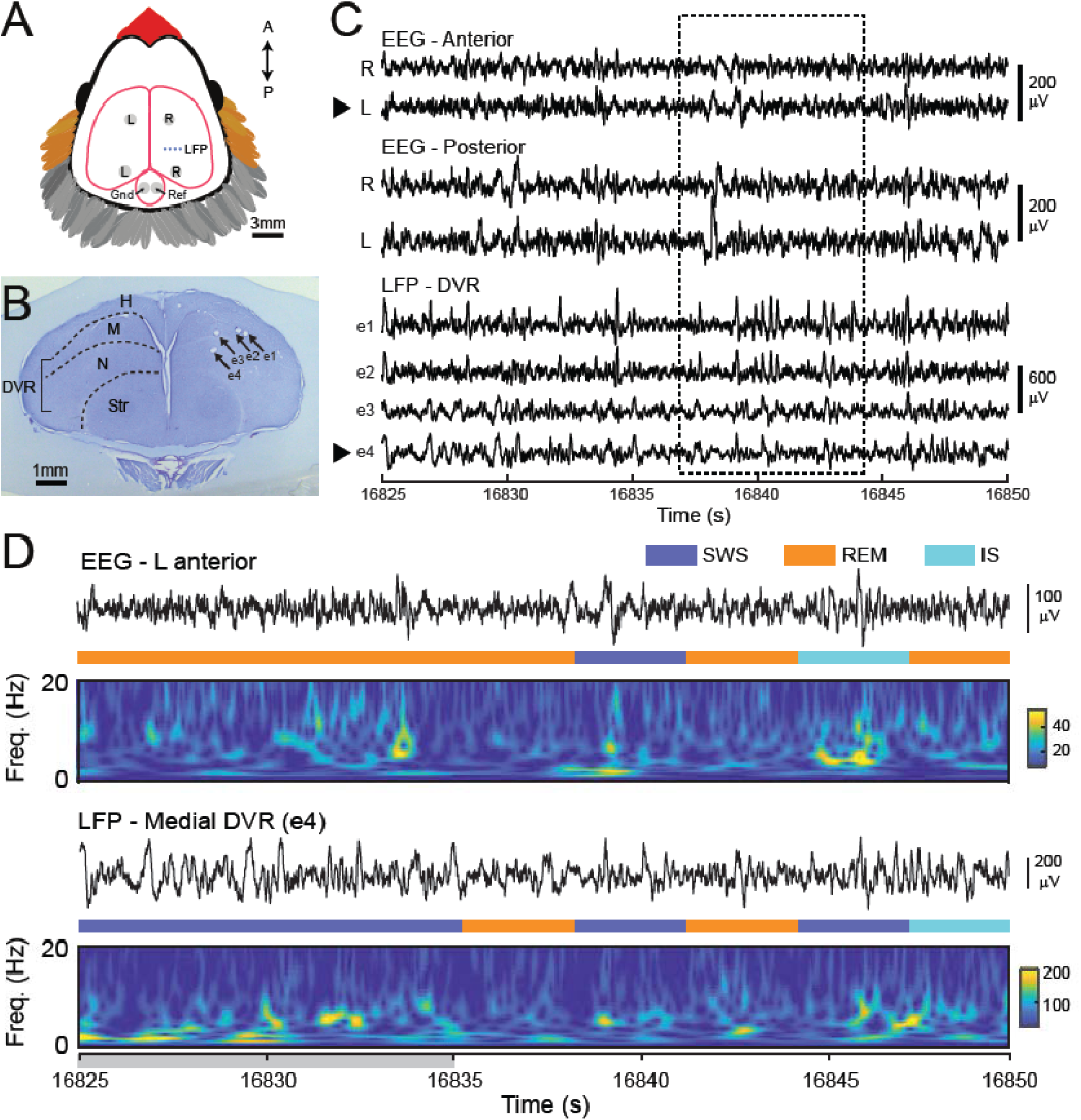
Local variations in sleep oscillations across pallial areas. **(A)** Schematic depicts the locations of the EEG and LFP electrodes. 2 silver-ball EEG electrodes were implanted above the hyperpalium in both the right (R) and left (L) hemisphere in an anterior (A) and posterior (P) position. 4-8 steel wire LFP electrodes were implanted in the right hemisphere in 3 birds. Of the 4 to 8 LFP electrodes, only the most medial LFP electrode (e.g., e4) was used in the analysis. Two silver-ball EEG electrodes were placed over the cerebellum and used as the ground electrode (Gnd) and reference electrode (Ref). **(B)** Histological section from bird w042 depicts LFP electrode tracts (black arrows; e1-e4). Electrode tips were 1.8-2.2 mm deep in the mesopallium and nidopallium of the dorsal ventricular ridge (DVR). H, hyperpallium; M, mesopallium; N, nidopallium; Str, striatum. **(C)** Simultaneous recordings across different sites and depths reveal variations in sleep oscillations across pallial areas, with notable differences between EEG and LFP electrodes (dashed box). LFP traces from top to bottom correspond to sites from most lateral (e1) to most medial (e4). Black arrowheads indicate the data traces enlarged in (D). **(D)** Simultaneously recorded left anterior EEG trace (top) and the most medial LFP trace (e4; bottom) with corresponding sleep state labels (colored bars) and spectrograms. Gray shaded area (16825-16835 s) indicates a region of contrasting sleep state labels; note the differences in EEG and LFP amplitude (black traces) as well as the spectral content (spectrograms). Data traces are the same as those in (C).

The LFP electrodes were positioned in a row orthogonal to the midline, where the LFP electrode closest to the midline was positioned 1.5 mm lateral from the midline, and the LFP electrode most lateral was positioned 2.5 mm lateral from the midline. LFP electrodes were 2-3 mm frontal to the junction between the frontal and parietal cranial bones. The two frontal EEG electrodes were positioned 2 mm lateral from the midline on each side and 5 mm anterior to the junction of frontal and parietal cranial bones. Posterior EEG electrodes were placed 2.5 mm lateral to the midline and 1 mm posterior to the frontal and parietal bones junction. Reference and ground electrodes (silver ball electrodes) were positioned over the cerebellum (Fig. 1A).

After the surgery, animals were released from the stereotaxic frame and administered an analgesic (Metamizole (100-150 mg/kg, i.m.). Animals were kept on the heating pad until signs of recovery from the anesthesia were observed. Following the surgery, an antibiotic (Baytril, 1025 mg/kg, i.m.) and an analgesic (Metamizole (100-150 mg/kg, i.m.) were administered for up to 3 days.

### Electrophysiology

After surgery, the birds were habituated to the implant within the recording chambers. During a recording session, the preamplifier (Intan Technologies, RHD2132) and its associated tether cable were connected to the electrode implant one hour before the lights were turned off (light-off period: 10:00 PM - 10:00 AM) to allow the birds to adopt a natural sleeping position. The weight of the tether cable was counterbalanced using an elastic rubber band. Electrode signals were acquired at a sampling rate of 30 kHz using an Open Ephys data acquisition board (OEPS, Portugal), filtered within the range of 0.1-9987 Hz, and saved in the Open Ephys GUI “continuous” format. A near-infrared sensitive camera (acA1300-60gm, Basler Ag, Germany) was used to acquire video recordings of the animals using custom-written software (ZR View, Robert Zollner). Camera frames were triggered using a pulse generator (Pulse Pal, Sanworks, NY, USA (35)) at a frame rate of 20 Hz. The tether cable was disconnected from the headstage half an hour after the lights were turned on the following morning.

### Histology

After the final recording session, the birds were deeply anesthetized using a lethal dose of sodium pentobarbital (250 mg/kg, i.m.). Once the corneal reflex and heartbeat had ceased, the bird was decapitated and the brain was extracted from the skull. The brain was subsequently immersed in a 4% paraformaldehyde solution in phosphate-buffered saline for a minimum duration of one day. 80 µm coronal sections were stained with cresyl violet to aid in electrode site verification.

### Data Analysis

Data were recorded from the birds for a period of approximately two weeks. The nighttime recordings analyzed in this study occurred after at least one week of habituation to the recording environment. For data analysis, one whole night of sleep (i.e., 12 hours) was analyzed for each bird. The nighttime recording was selected on the basis of behavior (i.e., sleeping behavior occurred in one long session and was not broken up during the night), the quality of electrophysiology data, and the quality of the video recording (the bird was visible within the recording frame throughout the night).

### Automatic sleep staging

All data analysis was conducted offline in Matlab (Matworks, Inc., Natick, MA). Electrophysiological data were down-sampled to 468.75 samples/s (equivalent to a down-sampling factor of 64). Similar to previous work (Yeganegi & Ondracek, 2023), a semi-automatic sleep staging procedure was employed to classify each 3-second data window into one of the following categories: awake, SWS, REM sleep, or Intermediate Sleep (IS). IS can be considered a transitional stage of NREM sleep, and it typically lacks the large slow oscillations that characterize SWS (Low et al., 2008).

In a first pass, the distinction between wake and sleep was determined by examining each 3-s bin for the strong presence of gamma activity (30-50 Hz), since the wake-associated body movements were reflected in artifacts within this band. For this detection, the moving median and the moving interquartile interval (IQR) were computed in 10-minute windows for the data from the left anterior EEG channel. The 3-sec bins where 30-50 Hz power exceeded the threshold of (moving median + 3×moving IQR) were considered “wake”. Wake detection was confirmed by inspecting the corresponding video frames; movement artifacts typically involved body twitches, eye-opening, or changes in posture.

Next, we segmented the rest of the bins, i.e., sleep bins, into three clusters of SWS, REM sleep, and IS by comparing the δ/γ power ratio in each bin. Similar to the detection of wake bins, the δ/γ in each 3-sec bin was compared to the moving median (δ/γ) + *A*×moving IQR(δ/γ) for SWS and moving median(δ/γ) - *A*×moving IQR(δ/γ) for REM sleep. The remaining bins were labeled as IS. Here *A* is a parameter that we iteratively increment from 0.1 to 2 to find the best clustering output. For each value of *A* we checked the output via two visualizations. 1) Visualization of the clustering results in a 3D space (Supplementary Fig. S2) composed of log(δ), log(γ), and max amplitude in each bin, similar to (Low et al., 2008) and (Canavan & Margoliash, 2020). 2) Visualization of the labels alongside the raw EEG trace (Supplementary Fig. S2). Based on these two visualizations, we selected the value of *A* with which the clustering output was maximally crisp and accepted the corresponding clustering results.

### Sleep state congruence

To quantify differences in neural activity across different brain regions, we first scored the sleep states separately in each of the five electrode sites: right and left anterior EEG, right and left posterior EEG, and the medial-most LFP electrode in DVR. Then, we used the definition of congruence as stated in (Durán et al., 2018); that is, “congruence” was defined as the percentage with which a given electrode shows the same sleep stage as a baseline electrode. We chose the left anterior EEG electrode as the baseline electrode, similar to (Durán et al., 2018). Congruence was then computed for each pairwise electrode comparison for each stage of sleep.

For example, to determine the congruence between the right posterior electrode and the baseline electrode (left anterior EEG) during SWS, we identified all of the 3-s bins that were labeled as SWS in the baseline electrode, and then calculated the percentage of those bins that were also labeled as SWS in the right posterior electrode. If all SWS bins in the baseline electrode corresponded to matching SWS bins in the comparison electrode, then the congruence was 100%. If some of those bins did not match, i.e., a SWS bin in the baseline electrode was labeled as a IS bin in the comparison electrode, then the congruence measurement equaled less than 100%.

### Spectral analysis

Power spectral analysis was computed using the Welch method on 3-s windows of data for each electrode. The number of discrete Fourier transform (DFT) points to use in the PSD estimate was set to 512. A 95% confidence interval was then estimated by computing the standard error as described in (Altman & Bland, 2005). To estimate the spectrograms visualized in Fig. 1D, the continuous wavelet transform (‘cwt’ function in Matlab) was used with a Gabor kernel

### Comparison of δ/γ power ratio

As an alternative approach to the congruence analysis, we compared the δ/γ power ratio between the most medial LFP channel and the left anterior EEG channels, i.e., the same channels used in the congruence analysis. δ and γ were defined as spectral band power in 1-4 Hz and 30-48 Hz respectively. First, the δ/γ power ratio was calculated in each channel in 3-sec bins and then smoothed in 30-sec windows to factor out transient fluctuations in the power. In each bird, this resulted in two vectors of δ/γ values, one for the EEG and one for the LFP signals. Then, each δ/γ vector of values in each channel, EEG and LFP, were normalized by their range, so that both δ/γ vector of values in the EEG and LFP channel had the same range, that is, zero to one. Finally, we found the difference between the two δ/γ vectors by subtracting them in each bin and calculating the absolute value of the difference.

### Temporal characteristics of congruence

In order to track the temporal characteristics of congruence and specifically how it evolved over a night of sleep, we also computed the congruence in sliding windows throughout the night. The window length was set to five minutes so that we had enough bins of each stage of sleep for the congruence calculation while maintaining a high temporal resolution.

### Sleep stage transition analysis

To investigate possible temporal precedence between the left and right hemispheres during the transitions between REM sleep and SWS stages, we compared the spectrograms of the EEGs between the left and right hemispheres. These comparisons were performed separately for the anterior and posterior channels between the hemispheres. First, based on the sleep scoring labels in the left anterior EEG (used as reference in the congruence analysis), we found all periods of SWS longer than one bin which were followed immediately by a period of REM sleep longer than one bin (and vice versa for REM sleep to SWS). Then, we calculated the spectrograms around these transition times in the left and right hemisphere EEGs separately, using continuous wavelet transform with Gabor kernels. Finally, we calculated the sum of power in the 5-25 Hz band in the spectrograms to obtain the transition-locked spectral power traces.

### Statistical analysis

To compare the duration and the percentage of each stage of sleep across the recording sits, we used the Wilcoxon rank-sum test (also known as the Mann-Whitney U-test). The U-test is a non-parametric test which does not require the assumption of a normal-distribution. Since we aimed to make the comparisons between five different sites, that is, the four EEGs and the most medial LFP, we considered 10 pair-wise comparisons and used a post-hoc Bonferroni correction to determine significance (e.g., α = 0.05 == 0.005 after correction; α = 0.01 == 0.0001 after correction). Mean and standard errors are also reported (See Supplementary Information for statistical tests).

## Results

### Local variations in sleep state are present across the avian pallium

We recorded surface EEGs from the left and right hemisphere over the hyperpallium and LFPs from the DVR (nidopallium and mesopallium) in three zebra finches. Two silver ball electrodes recorded surface EEG activity in anterior and posterior sites in each hemisphere (four EEG electrodes in total) (Fig. 1A). In addition, 4-8 steel electrodes recorded the LFP in the DVR (Fig. 1A and 1B). Of the 4-8 LFP electrodes, only the LFP signal from the most medial electrode was used in the analysis (Fig 1B, electrode e4); the other LFP signals were not analyzed. During sleep, both EEG and LFP electrodes showed brain oscillations typical of sleep, i.e., alternative periods of high amplitude slow wave activity, periods of low amplitude gamma oscillations, and periods of mixed frequency EEG activity. A 25-sec trace of EEG and LFP activity during sleep is depicted in Fig. 1C.

As a whole, the EEG and LFP activity was highly heterogeneous across sites (Fig. 1C, dashed box): high-amplitude peaks, deflections, and bursts of activity appeared on only one EEG electrode or only in the LFP recordings. In addition to these fast variations, we also observed sustained periods of time when different oscillations appeared simultaneously across different sites (Fig. 1D, red shading). To better characterize these local differences in neural activity, we segmented the EEG and LFP into discrete sleep states for each electrode separately using a sleep segmentation algorithm that clustered and scored the sleep data on the basis of spectral features typical of avian sleep (Canavan & Margoliash, 2020; Low et al., 2008). We labeled each 3-sec bin of neural activity as either of awake, SWS, IS (intermediate sleep), or REM sleep (Fig. 1D). The differences in sleep states, as scored individually across different electrode sites, correspond with the local variations in the spectral composition of the neural activity at these sites (gray shading, Fig. 1D).

### Sleep stage segmentation are in line with previously published results

We first quantified the statistics of the sleep staging as a function of the recording sites. We compared the overall percentage of time spent in each sleep state for each electrode site (Fig. 2A). Percentages of time spent in SWS, REM sleep, and IS sleep states were not significantly different as a function of electrode site (Mann-Whitney U-test, Bonferroni correction for multiple comparisons, p > 0.005 for all comparisons).

**Fig. 2.**
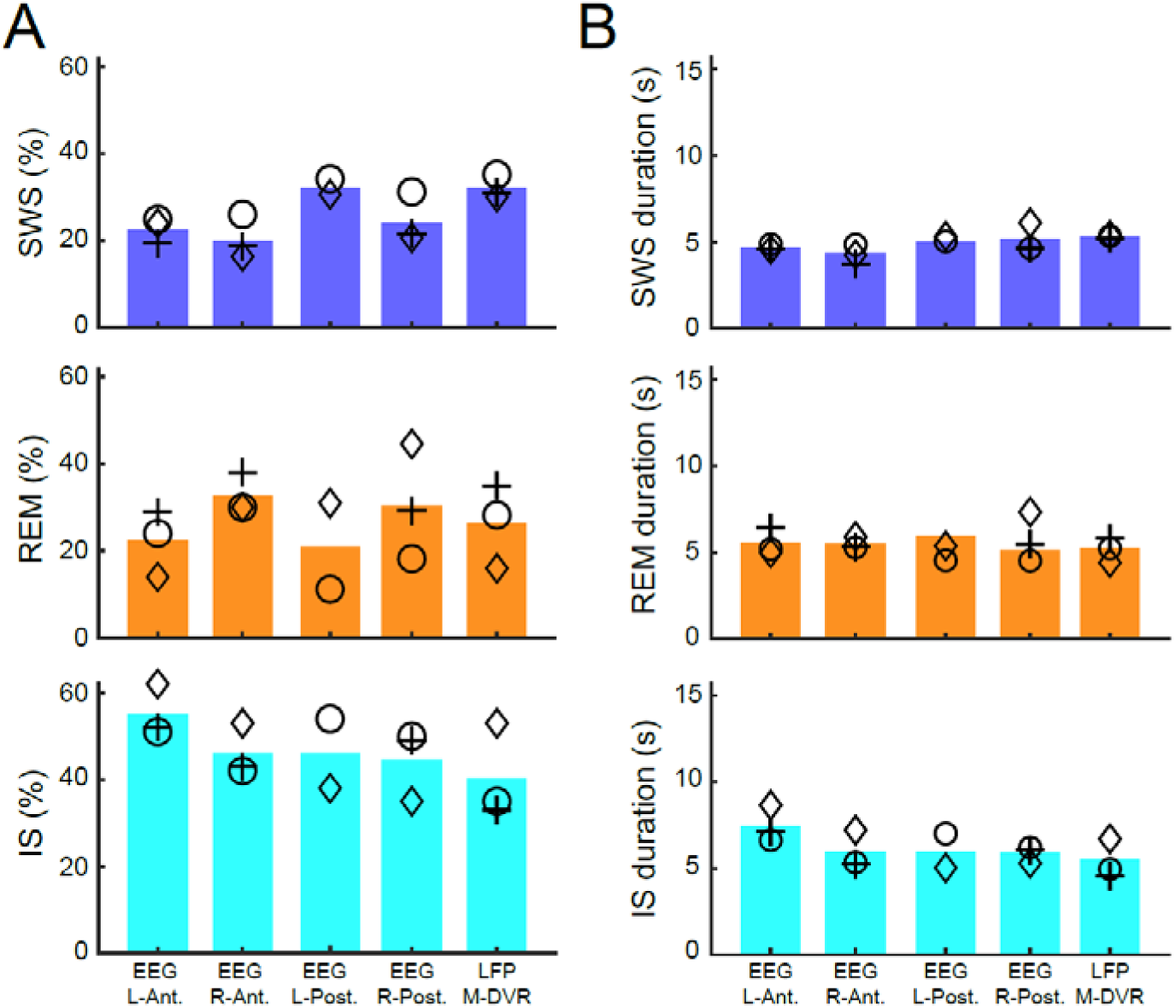
Sleep stage durations vary as a function of electrode location. **(A)** Bar plots indicate the mean percentages of SWS (top), REM sleep (middle), and IS (bottom) pooled over all nights (n=3) for the 5 different recording sites. Symbols indicate individual animals (see Table 1 for details). **(B)** Mean durations for SWS, REM sleep, and IS sleep states. Figure conventions same as in (A). Notably, sleep state durations were significantly different as a function of recording location, particularly for SWS and IS stages. Although the differences in the duration across nights are small, in most of the cases, they were statistically significant (see the Supplementary Information for the statistical tests).

We also analyzed the duration of time spent in SWS, IS, and REM sleep across the different recording sites. Bouts of SWS were significantly longer for the LFP electrode by around 1 sec compared to the EEG sites (Fig. 2B; see Supplementary Information for statistical tests), but overall the differences between electrodes were minimal. These results are in line with previously published results (Yeganegi & Ondracek, 2023).

### Congruence analysis reveals local sleep states

Next we asked how frequently two distant electrode sites share the same sleep stage at the same time. Our data suggested that this was a relatively rare event: we observed that sleep scores were infrequently identical for distinct recording sites (Fig. 3A, B). In order to quantify this, we calculated the “congruence” between each electrode and a designated baseline electrode (Durán et al., 2018)). Matching (Durán et al., 2018), we chose to use the left anterior EEG electrode as the baseline electrode (Fig. 3A, B). Congruence is an estimate of the probability that two electrode sites display the same sleep state at the same time (see Methods for details). Congruence values less than 100% indicate a state difference between the two electrodes sites and thus identify local differences in sleep states. The congruence values for each electrode and for each sleep state are presented in Fig. 3C. We observed that for all sleep states and across all electrodes, the congruence did not surpass 54%. This suggests that the neural oscillations during sleep, to a large extent, occur within a small anatomical network. In other words, a given sleep state is not a brain-wide phenomenon, but is a local event.

**Fig. 3.**
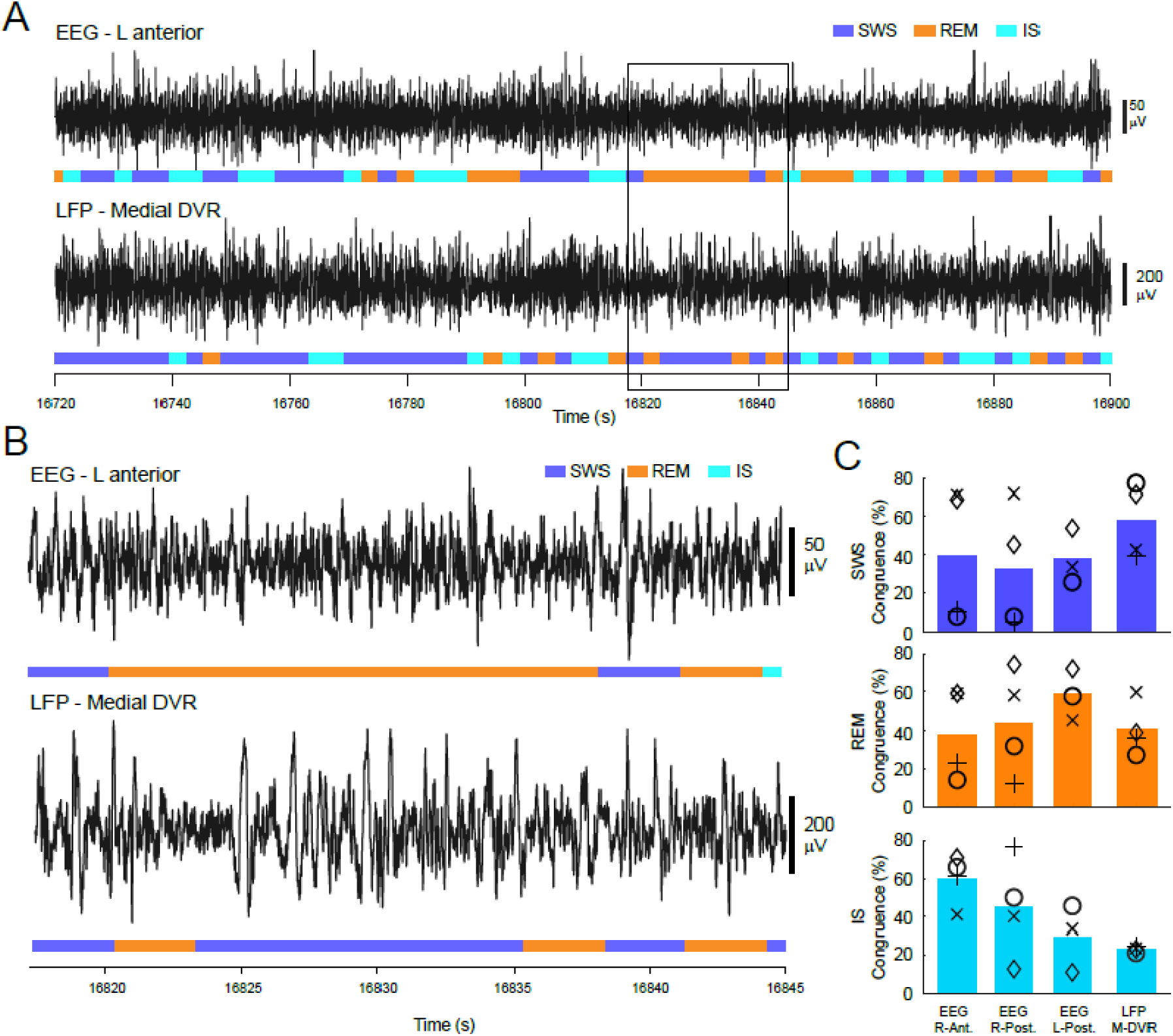
Quantification of simultaneous local differences in sleep state across the recording sites. **(A)** Traces depict the simultaneous left anterior EEG and medial DVR LFP with the corresponding sleep states scores (colored bars). Black box indicates a section that is enlarged in **(B).** Sleep state differences between the recording sites are evident through the color-coded representations of sleep stages for the same time windows. **(C)** Bar plots indicate the mean percentages of congruence for SWS (top), REM sleep (middle), and IS (bottom) pooled over all nights (n=3) for the 4 different recording sites compared to the baseline electrode (the left anterior EEG). In all cases, the mean congruence was below 54.0%. The lowest mean congruence (23.2%) was observed in the DVR during IS.

### δ/γ power of neural activity differs in DVR and hyperpallium

Critically, the congruence analysis outlined in the preceding section may be influenced by the distinct outputs of the clustering algorithm when applied to EEG and LFP signals. To address this potential confound and provide an alternative assessment of inter-site differences, we compared the δ/γ power ratio between the most medial LFP and the left anterior EEG (the baseline electrode in congruence analysis). Although the δ/γ power ratio of the EEG and LFP fluctuate generally in conjunction, we frequently observed periods in which the two δ/γ power ratios diverge from each other (Fig. 4A). For a considerable portion of bins (26.0±1.7%, averaged across birds), the difference between the two δ/γ ratios was above 10% (see the example in Fig. 4B, and the histograms in Fig. 4D). Importantly, these periods of divergence of δ/γ ratios overlaps with the time bins where sleep scoring labels in EEG and LFP are different (in Fig. 4, juxtapose B and C). This analysis further supplements our argument of dissimilar sleep stages that appear simultaneously across the pallial sites.

**Fig. 4.**
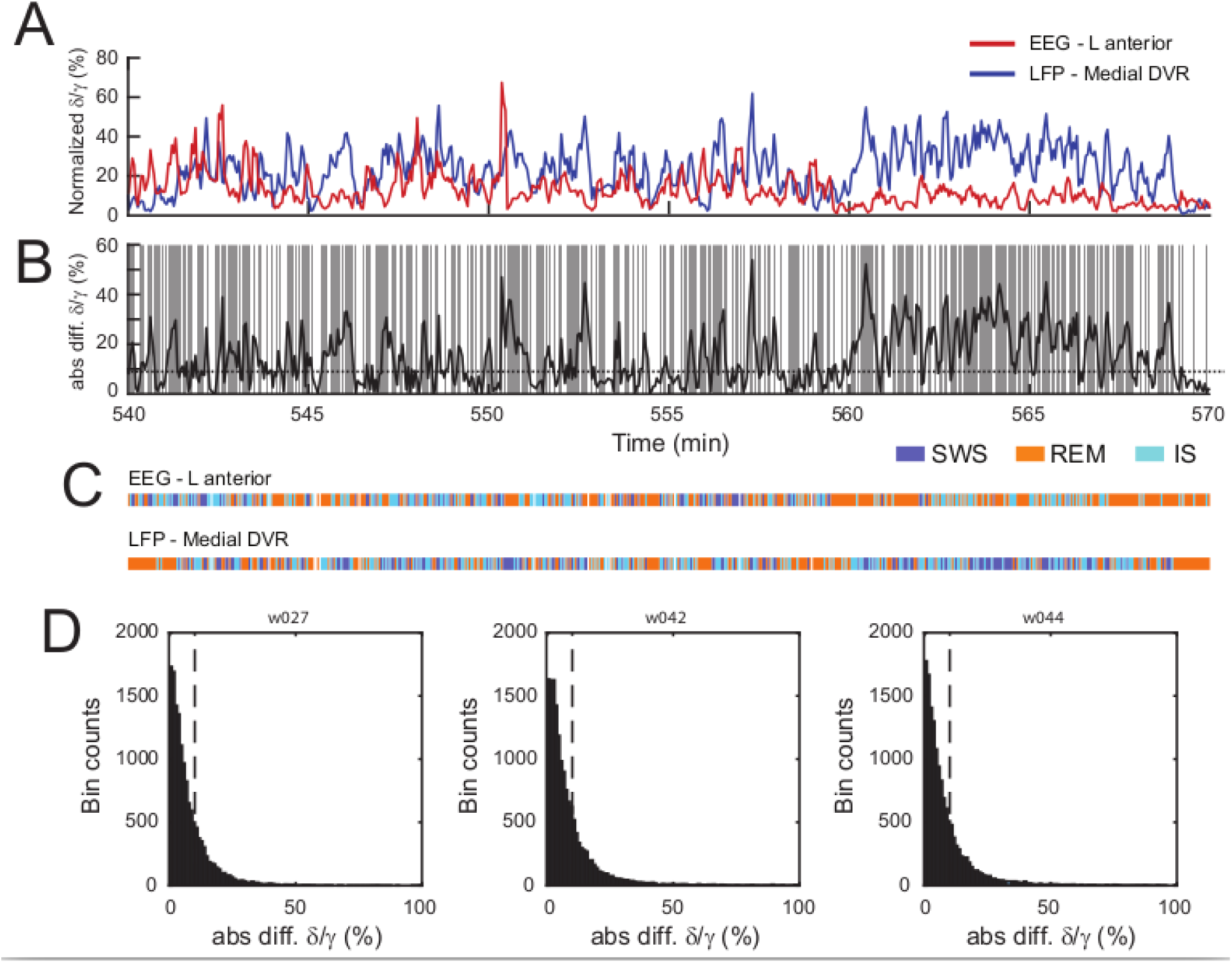
δ/γ power ratios show divergence between EEG and LFP signals. **(A)** Traces show the normalized δ/γ power ratio for the left anterior EEG baseline electrode (red line) and the most medial LFP electrode (blue line). Note the divergence between the traces during the last ten minutes. δ/γ normalization for visualization purposes only. **(B)** The absolute value of the difference between the EEG and LFP δ/γ ratios. Dashed line indicates the 10% difference threshold. Gray shading indicates divergence values that exceeded 10%. **(C)** Corresponding sleep scoring labels for the EEG and LFP for the same period as in (A) and (B). Note that periods with dissimilar sleep stage labels in the EEG and LFP (gray shading in B) coincide with periods of high divergence of δ/γ ratios in (B). **(D)** Histograms illustrate the distribution of δ/γ ratio differences between the EEG and LFP. For a large portion of bins, the divergence was above 10% (10% threshold indicated with dashed line; exact percentage of bins above 10%: 24%, 27%, and 27% from left to right).

### Congruence of sleep states changes throughout the night

To explore the temporal dynamics of local sleep as it evolved during sleep, we also tracked the changes in the “instantaneous” congruence between the recording sites by computing the congruence in 5-minute long sliding windows (Fig. 5, A and B). We focused this analysis on the DVR electrode and the left anterior EEG. We observed that the instantaneous congruence between the right DVR and the left hyperpallium did indeed change throughout the sleep time and was largely dependent on the sleep state.

**Fig. 5.**
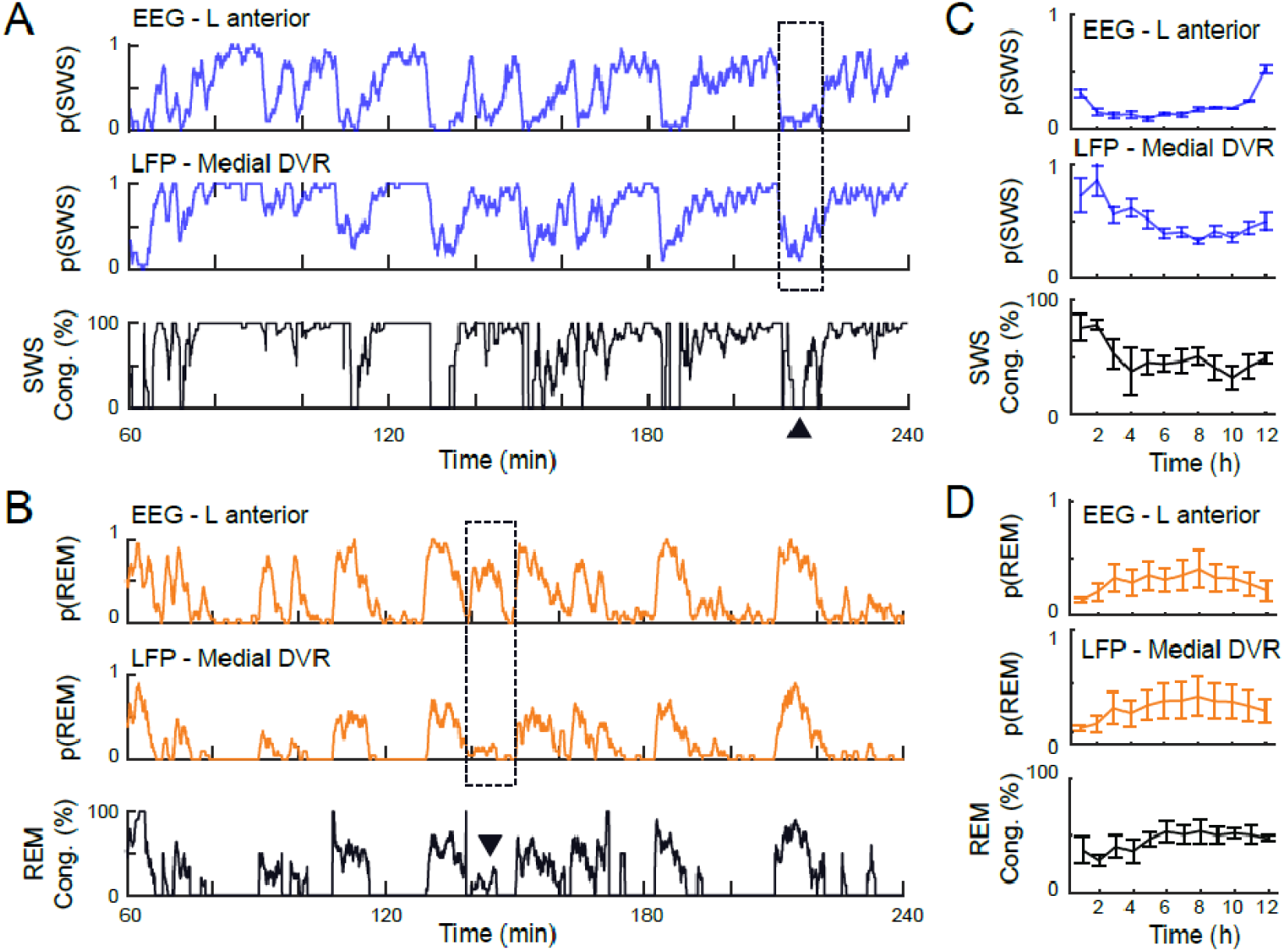
Instantaneous congruence changes throughout the night. **(A)** Instantaneous likelihood of SWS, p(SWS), in the baseline electrode (top), in the DVR (middle), and the congruence of SWS between these two sites (bottom) are shown. Congruence dynamics were closely tied to the likelihood of SWS in both channels, especially the DVR channel. The dotted box and arrowhead in the lower subplot highlights a period of low congruence between the two electrodes. This period coincides with a differential likelihood of SWS in the DVR and the left anterior EEG. **(B)** Similar to (A), but for the dynamics of the REM sleep. Note again that the marked period with low REM sleep congruence coincides with different likelihood of REM sleep in the two recording sites. **(C)** SWS shows a characteristic decrease for the first half of sleep time, both in the left anterior EEG and in the DVR. The congruence between the DVR and the left anterior EEG changes during sleep time in concert with the likelihood of SWS in the DVR. **(D)** In contrast to SWS, REM sleep likelihood increases both in the left anterior EEG and in the DVR for the first half of sleep time. Similar to SWS, the congruence of REM sleep episodes between the DVR and the left anterior EEG changes during sleep time in concert with the likelihood of REM sleep in the DVR.

During the first half of night, the congruence between the DVR and baseline electrode decreased for SWS and increased for REM sleep (Fig. 5, C and D). This temporal dynamic was closely coupled to the temporal dynamics of SWS and REM sleep in DVR, and not in the baseline channel. To prove this, we calculated the correlation coefficient between the instantaneous congruence and the likelihood of the corresponding stage of sleep, for both SWS and REM sleep stages (the instantaneous likelihood of SWS and REM sleep are denoted by p(SWS) and p(REM) in Fig. 5). The instantaneous congruence of SWS in the DVR was highly correlated with the likelihood of SWS in the DVR: 0.86 ± 0.12 (correlation coefficient, mean and std among birds). Similar results were true for the REM sleep state (0.93 ± 0.04). In contrast, the correlation coefficient between the congruence and the likelihood of SWS and REM sleep in the baseline electrode (anterior left EEG) was comparably low: 0.47 ± 0.18 for SWS (mean and sd among birds) and 0.52 ± 0.31 for the REM sleep state.

### Transient hemispheric asymmetries during state transitions

Lastly, we analyzed whether consistent hemispheric differences exist in the timing of sleep state transitions (that is, transitions from SWS to REM sleep and vice versa). To that end, we compared the transition-locked spectrograms of the left and right EEG channels (Fig. 6, top two rows). Our analysis revealed that across both hemispheres, power in the 5-25 Hz frequency band increased during transitions to SWS and decreased during transitions to REM sleep. Aggregated power within this frequency band was computed and analyzed (Fig. 6, bottom row). The spectral profiles from left and right hemispheres showed a high degree of similarity, with no clear temporal precedence observed at the temporal scale we examined. However, within these similar patterns, transient divergences in power profiles were detected. Particularly, ∼300 ms before REM-to-SWS transitions, a slight differential increase in power was observed in the left hemisphere compared to the right (indicated by arrows in Fig. 6B and 6D). However, this difference did not reach statistical significance (rank-sum test; p > 0.05). In summary, our data suggests the presence of asymmetric activity between the hemispheres during sleep state transitions, but these differences were not statistically robust.

**Fig 6.**
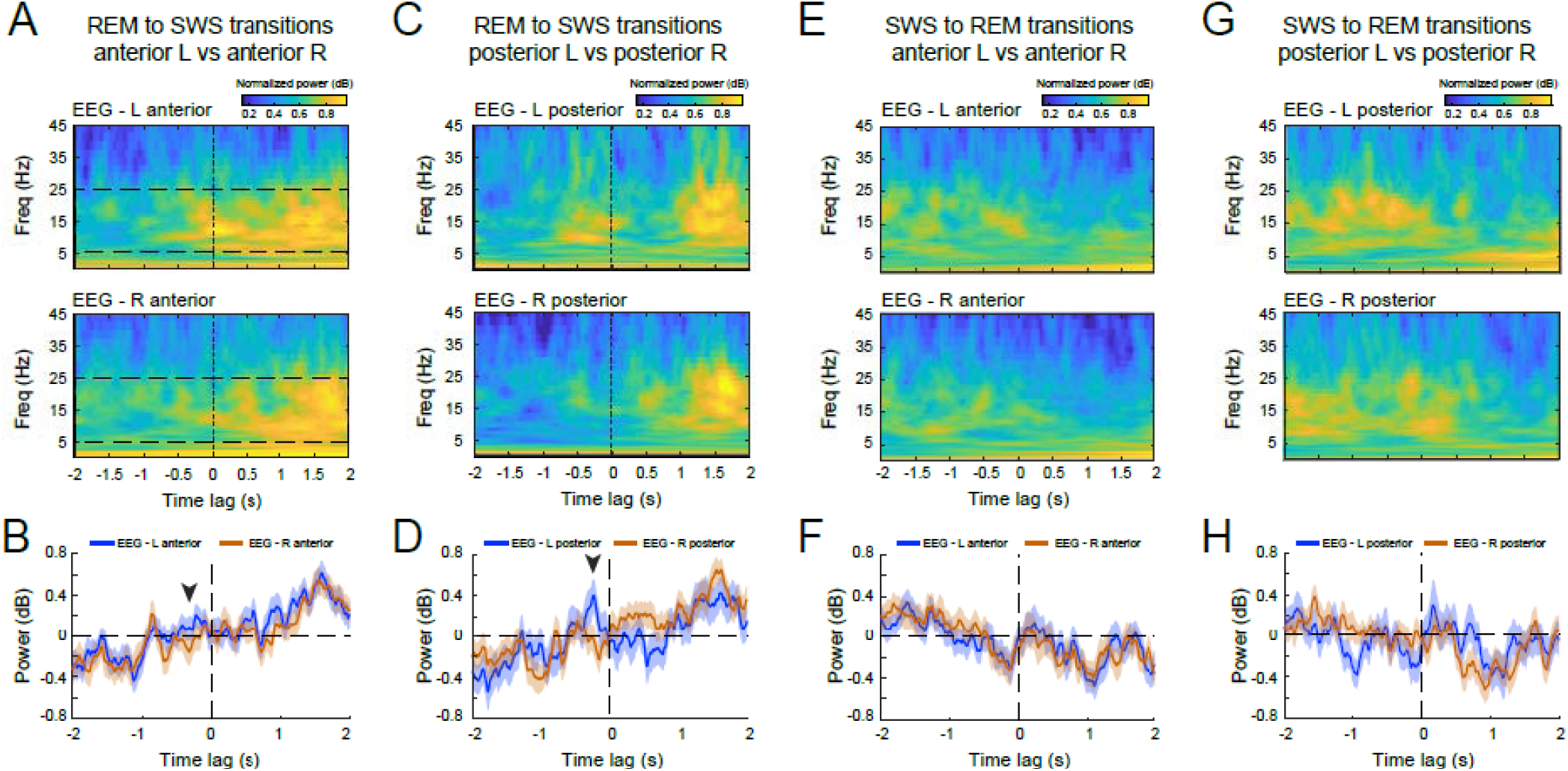
Spectral analysis of sleep state transitions across hemispheres. **(A)** Spectrograms indicate the peri-transitions time window for REM-to-SWS transitions for the anterior left (top) and anterior right (bottom) EEG channels. **(B)** The aggregated spectral power in the 5-25 Hz band for EEG data locked to REM-to-SWS transitions are displayed for the anterior left EEG channel (blue line) and anterior right EEG channel (red line). Black arrowhead indicates a transient divergence between the two electrodes at approximately 300 ms before the REM-to-SWS; however, this hemispheric difference was not statistically significant. **(C, D)** Figure conventions same as in (A, B); data displayed for posterior left and posterior right EEG electrodes during REM-to-SWS transitions. In (D), note similar divergence between electrode sites as in (B; black arrowhead). **(E)**. Data displayed for the anterior left (top) and anterior right (bottom) EEG channels for SWS-REM transitions. **(F)** The aggregated spectral power for EEG data locked to SWS-to-REM transitions are displayed for the anterior left EEG channel (blue line) and anterior right EEG channel (red line). Note the tight correspondence between traces, in contrast to the REM-SWS transitions (e.g., B, D). **(G, H)** Figure conventions same as in (E, F); data displayed for posterior left and posterior right EEG electrodes during SWS-to-REM transitions.

## Discussion

Our findings highlight the presence of local neural oscillations linked to discrete sleep stages across diverse sites within the avian pallium, which is in line with emerging evidence that suggests that sleep may be locally regulated across the brain (Krueger et al., 2008).

Local regulation of sleep under homeostatic control has already been demonstrated in birds. In (Lesku et al., 2011), one eye in the pigeons was covered, while the other eye was presented with a visual stimulus. On the following night, after the asymmetric visual exposure, EEG was recorded during sleep from the left and right hemispheres. The slow wave activity increased only in the hemisphere contralateral to the visually stimulated eye, demonstrating the local regulation of sleep as a homeostatic mechanism. In contrast to these results, our findings show that local differences in sleep also exist after unperturbed awake activity and is not necessarily the result of homeostatic control.

The idea that sleep could be regulated locally challenges conventional theories that favor the concept of brain-wide sleep that is driven by sleep regulatory *centers* (Saper et al., 2010; Scammell et al., 2017). Indeed, recent studies have suggested that sleep may be a default characteristic of neuronal networks, as demonstrated by the observation of sleep-like oscillations in isolated neural tissues (Bandarabadi et al., 2020; Hinard et al., 2012) and relies on preceding (asymmetrical) activity during wakefulness (Huber et al., 2004; Kattler et al., 1994; Vyazovskiy et al., 2000). Other studies have suggested that metabolically driven changes in sleep-regulatory substances could also regulate local sleep patterns (Hinard et al., 2012). Finally, mathematical modeling has highlighted that the synchronization of sleep-like states across individual cortical columns could ultimately culminate in brain-wide sleep as an emergent property of local-network interactions, without a need for a central switch (Roy et al., 2008). These recent findings collectively propose that the regulation of sleep might not exclusively hinge on a designated neural circuit that switches whole cortical areas between discrete states. Our observation of distinct sleep stages existing simultaneously across sites within the avian pallium adds evidence to this growing body of emerging data.

The original work that quantified the homogeneity of sleep stages across the cortical sites using congruence was conducted in rats (Durán et al., 2018). In that study, the activity in cortical sites (medial prefrontal, and superficial parietal) and a hippocampal site was compared to a baseline cortical site (superficial frontal). The authors reported average congruence during SWS and REM sleep above 87%. But during IS sleep, the value dropped to 36%. In comparison, in the zebra finch pallium, we observed that the average congruence over all three stages was not higher than 54%, without a significant distinction between the IS and the SWS and REM sleep states (see Fig. 3C). These findings were further supported by our comparison of the δ/γ power ratio for the LFP and left anterior EEG channel (Fig. 4). Here we saw that for as much as 27 % of the sleep time, the δ/γ power ratios calculated individually for these electrodes deviated by more than 10%. As such, we have found a much higher level of heterogeneity in the sleep stages across avian pallium as compared to the mammalian brain.

This observation could be explained considering the differences in the pallial designs between the mammalian and avian brains, most prominently the laminar organization of cortical areas and the presence of the corpus callosum in the placental mammal’s brain. However, there are caveats to this explanation. First, the technical differences in sleep scoring methods could account for differences in the estimated congruence. Second, in rats, the IS sleep occupied only 2.2% percentage of the time, including the wake period. In this work (Fig. 2A), we observed that the IS sleep occupies a much larger percentage of sleep (more than 40%) in all channels. Third, the differences in the recording sites might account for marked differences. For instance, it has been reported that under isoflurane anesthesia, slow waves are present in the nidopallium (a subdivision of avian pallium ventral to mesopallium) but not the avian hippocampus (van der Meij et al., 2020).

Finally, our data has implications regarding the origin of slow waves in the avian pallium. In the mammalian brain, slow oscillations seem to emerge from various neocortical regions, subsequently propagating horizontally across the neocortex akin to a traveling wave (Murphy et al., 2009; Nir et al., 2011). Although the neocortex possesses the inherent capacity to generate slow oscillations after recovery from thalamotomy, thalamic input plays a critical role in the generation of slow waves under normal physiological circumstances (Crunelli & Hughes, 2010; Lemieux et al., 2014). Notably, the inception of slow waves tends to manifest predominantly within layer 5, a layer that receives inputs from the thalamus (Constantinople & Bruno, 2013), and subsequently propagates vertically within cortical columns (Capone et al., 2019).

In the avian brain, two subregions of the visual hyperpallium are known to receive extensive thalamic input, namely the Interstitial part of the apical hyperpallium and the intercalated part of the hyperpallium (Reiner et al., 2004; Wild, 1987). A multichannel LFP recording that allowed simultaneous recordings from these two regions as well as a subregion with far less thalamic input, i.e., hyperpallium densocellulare, showed that the amplitude of slow waves was higher in the thalamic recipient subdivisions (van der Meij et al., 2019). In addition, it was reported that the slow waves tend to appear first in these subdivisions and then propagate through, and outward, from these areas. Slow wave amplitude during sleep has not been directly investigated in the songbird pallium, which also receives extensive thalamic input to brain areas involved in vocal learning (Akutagawa & Konishi, 2005; Boettiger & Doupe, 1998; Coleman et al., 2007; Ondracek & Hahnloser, 2013).

Could local EEG patterns be regulated by thalamic input, or is it instead locally regulated by the pallium itself? Given the distinctions between the mammalian cortex and the avian pallium, further investigations are necessary to elucidate the underlying mechanisms responsible for the genesis of slow waves and their local modulations in the avian brain.

## Author Contributions

Conceptualization, H.Y. and J.M.O.; Methodology, H.Y. and J.M.O.; Software: H.Y. and J.M.O.; Validation, J.M.O.; Formal Analysis, H.Y.; Investigation, H.Y.; Writing – Original Draft, H.Y.; Writing – Review & Editing, J.M.O.; Funding Acquisition, J.M.O.; Resources, J.M.O.; Visualization, H.Y. and J.M.O.; Supervision, J.M.O.; Project Administration, J.M.O.

## Acknowledgements

The authors are thankful to B. Seibel and Y. Schwarz for technical assistance; C. Fink and E. Jochen for help with mechanical design, fabrication, and electronics; A. Schuhbauer for administrative assistance, and H. Luksch, for feedback and support of this research.

## Funding Information

This study was funded by grants from the German Research Foundation (Deutsche Forschungsgemeinschaft; ON 151/1-1) and the Daimler and Benz Foundation (Postdoctoral scholarship) awarded to Janie M. Ondracek.

## Conflict of Interest Statement

The authors declare no conflicts of interest related to this work.

## Data Availability Statement

The data that support the findings will be available in upon request from the corresponding author.

## Notes

### Competing Interest Statement

The authors have declared no competing interest.

### Summary of Updates

1) we have extensively rewritten the introduction to provide better clarity n the topic of local sleep. 2) We have extensively redone the analysis after excluding one bird in which the LFP electrode was implanted in the left hemisphere instead of the right hemisphere. The current number of birds analyzed is n=3. 3) We have extended the analysis to include two new figures, Fig. 4 and Fig. 6. Fig. 4 provides visualizations for analysis that complements the congruence analysis. Fig. 6 provides visualizations for a new analysis examining spectral features across SWS to REM and REM to SWS transitions.

