## Supplementary Figures and Information for "Local sleep in songbirds: Different simultaneous sleep states across the avian pallium"

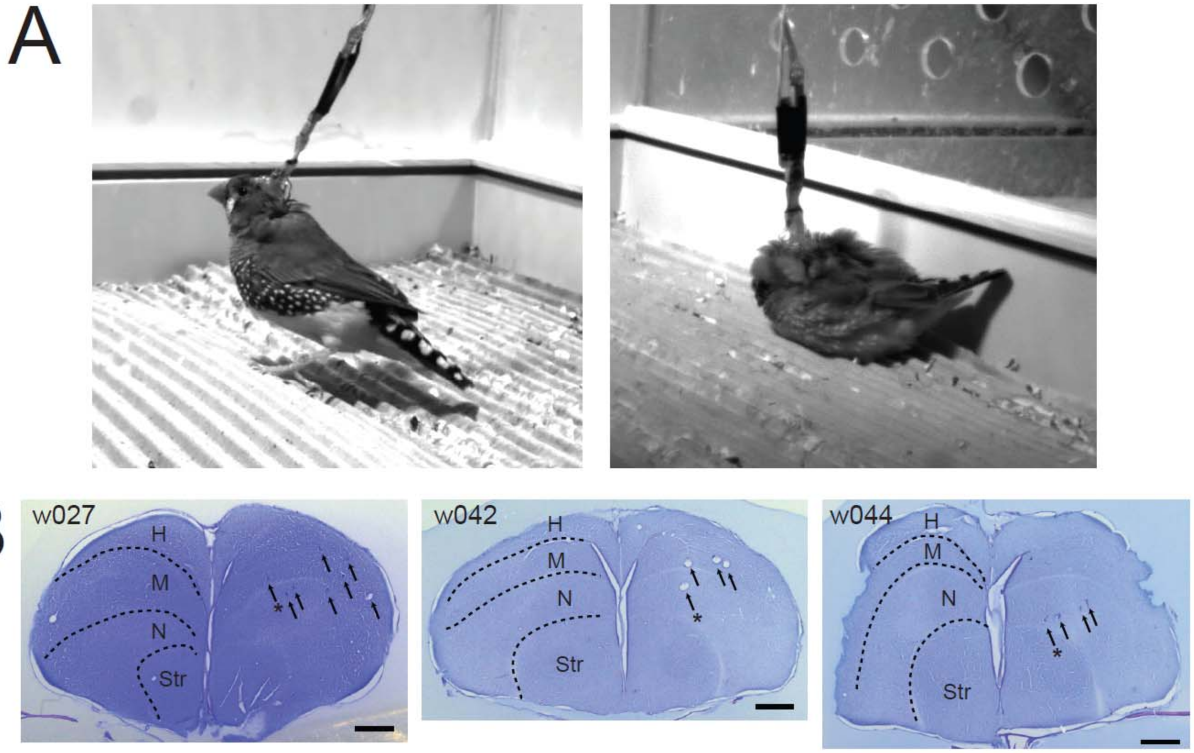


**Fig. S1 Animal posture and histology.** (A) Left, image of an awake tethered juvenile bird (w042). Right, image of the same bird sleeping. Note how bird is able to assume a sleep-specific posture despite the tether cable. (B) Histological sections stained with cresyl violet indicate the LFP electrode tracts (black arrows, n=8 LFP tracts in w027, n=4 LFP tracts in w042 and w044). Asterisk indicates the most medial LFP electrode that was used in the LFP analysis. Error bar indicates 1 mm. H, hyperpallium; M, mesopallium; N, nidopallium; Str, striatum.

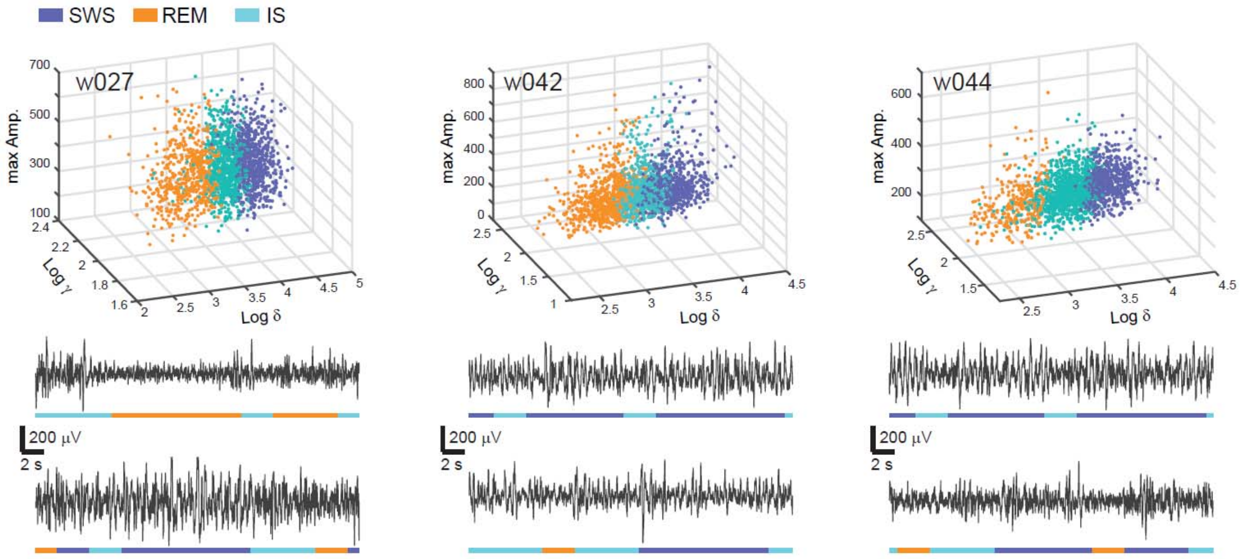


**Fig. S2 Examples of sleep staging for LFP signals.** Each panel (w027, left; w042, middle; and w044, right) is a 3-D plot of the 3-s bins of sleep time that were clustered as either SWS (blue), IS (cyan), or REM sleep (orange) for each bird. Each dot represents a different 3-s bin of sleep time, color coded according to its sleep state label. Note how sleep state clusters are segregated in 3-D spaces as a function of the spectral power of gamma (γ), delta (δ) and the maximum amplitude of the LFP signal. Below each panel are two 30-s long examples of the color-coded sleep state classification for the LFP channel for each bird. Note how orange REM states correspond to low amplitude LFP signals, blue SWS sleep states correspond to large-amplitude LFP signals, and cyan IS corresponds to an intermediate state.

### Supplementary Information

**Statistical Test Results**
**Differences in the Duration and Percentage of Sleep Stages Across Sites**

5 sites, so we will have 5*4/2=10 pair-wise tests.

The Wilcoxon rank sum test is used (a.k.a. Mann-Whitney U-test), because U-test is a non-parametric one, and dos not require normal-distribution assumption. For the multiple comparisons, i.e. 10 pair-wise tests, a Bonferroni correction factor (of 10) is applied. The mean and the standard error (SE=SD/sqrt(n)) is also reported.

α = 0.05 == 0.005
α = 0.01 == 0.001
α = 0.001 == 0.001

#### 1. Duration

**Test results for SWS:**
5 recording sites, which leads to 5*4/2=10 pair-wise tests.

The Wilcoxon ranksum test is used (a.k.a. Mann-Whitney U-test). The U-test is a non-parametric test, and does not require normal-distribution assumption. To account for the multiple comparison, i.e. 10 pair-wise comparisons, a Bonferroni correction factor of 10 is applied.
Mean and the standard error (SE=SD/sqrt(n)) are also reported.

**Test results for SWS:**
Between EEG-L-Ant. and EEG-R-Ant., p= 0.15906
N_EEG-L-Ant. = 6361, N_EEG-R-Ant. = 5276
mean: EEG-L-Ant. = 4.4455,  SE: EEG-L-Ant. = 0.040837
mean: EEG-R-Ant. = 4.3505,  SE: EEG-R-Ant. = 0.04225
-------
Between EEG-L-Ant. and EEG-R-Post., p= 9.1636e-05
N_EEG-L-Ant. = 6361, N_EEG-R-Post. = 5562
mean: EEG-L-Ant. = 4.4455,  SE: EEG-L-Ant. = 0.040837
mean: EEG-R-Post. = 4.938,  SE: EEG-R-Post. = 0.10873
-------
Between EEG-L-Ant. and EEG-L-Post., p= 4.6472e-15
N_EEG-L-Ant. = 6361, N_EEG-L-Post. = 6843
mean: EEG-L-Ant. = 4.4455,  SE: EEG-L-Ant. = 0.040837
mean: EEG-L-Post. = 4.9448,  SE: EEG-L-Post. = 0.054698
-------
Between EEG-L-Ant. and LFP, p= 3.2229e-51
N_EEG-L-Ant. = 6361, N_LFP = 8057
mean: EEG-L-Ant. = 4.4455,  SE: EEG-L-Ant. = 0.040837
mean: LFP = 5.4921,  SE: LFP = 0.057616
-------
Between EEG-R-Ant. and EEG-R-Post., p= 3.3184e-07
N_EEG-R-Ant. = 5276, N_EEG-R-Post. = 5562
mean: EEG-R-Ant. = 4.3505,  SE: EEG-R-Ant. = 0.04225
mean: EEG-R-Post. = 4.938,  SE: EEG-R-Post. = 0.10873
-------
Between EEG-R-Ant. and EEG-L-Post., p= 7.0596e-19

N_EEG-R-Ant. = 5276, N_EEG-L-Post. = 6843
mean: EEG-R-Ant. = 4.3505,  SE: EEG-R-Ant. = 0.04225
mean: EEG-L-Post. = 4.9448,  SE: EEG-L-Post. = 0.054698
-------
Between EEG-R-Ant. and LFP, p= 3.249e-55
N_EEG-R-Ant. = 5276, N_LFP = 8057
mean: EEG-R-Ant. = 4.3505,  SE: EEG-R-Ant. = 0.04225
mean: LFP = 5.4921,  SE: LFP = 0.057616
-------
Between EEG-R-Post. and EEG-L-Post., p= 0.00034267
N_EEG-R-Post. = 5562, N_EEG-L-Post. = 6843
mean: EEG-R-Post. = 4.938,  SE: EEG-R-Post. = 0.10873
mean: EEG-L-Post. = 4.9448,  SE: EEG-L-Post. = 0.054698
-------
Between EEG-R-Post. and LFP, p= 1.4066e-25
N_EEG-R-Post. = 5562, N_LFP = 8057
mean: EEG-R-Post. = 4.938,  SE: EEG-R-Post. = 0.10873
mean: LFP = 5.4921,  SE: LFP = 0.057616
-------
Between EEG-L-Post. and LFP, p= 2.6066e-13
N_EEG-L-Post. = 6843, N_LFP = 8057
mean: EEG-L-Post. = 4.9448,  SE: EEG-L-Post. = 0.054698
mean: LFP = 5.4921,  SE: LFP = 0.057616

**<u>Test results for REM:</u>**
Between EEG-L-Ant. and EEG-R-Ant., p= 2.1039e-37
N_EEG-L-Ant. = 5519, N_EEG-R-Ant. = 8103
mean: EEG-L-Ant. = 4.8579,  SE: EEG-L-Ant. = 0.054642
mean: EEG-R-Ant. = 5.8686,  SE: EEG-R-Ant. = 0.06276
-------
Between EEG-L-Ant. and EEG-R-Post., p= 2.2611e-46
N_EEG-L-Ant. = 5519, N_EEG-R-Post. = 7264
mean: EEG-L-Ant. = 4.8579,  SE: EEG-L-Ant. = 0.054642
mean: EEG-R-Post. = 6.235,  SE: EEG-R-Post. = 0.075723
-------
Between EEG-L-Ant. and EEG-L-Post., p= 2.5338e-35
N_EEG-L-Ant. = 5519, N_EEG-L-Post. = 6822
mean: EEG-L-Ant. = 4.8579,  SE: EEG-L-Ant. = 0.054642
mean: EEG-L-Post. = 5.8817,  SE: EEG-L-Post. = 0.066523
-------
Between EEG-L-Ant. and LFP, p= 2.4203e-14
N_EEG-L-Ant. = 5519, N_LFP = 6563
mean: EEG-L-Ant. = 4.8579,  SE: EEG-L-Ant. = 0.054642
mean: LFP = 5.473,  SE: LFP = 0.061067
-------
Between EEG-R-Ant. and EEG-R-Post., p= 0.030357
N_EEG-R-Ant. = 8103, N_EEG-R-Post. = 7264
mean: EEG-R-Ant. = 5.8686,  SE: EEG-R-Ant. = 0.06276
mean: EEG-R-Post. = 6.235,  SE: EEG-R-Post. = 0.075723
-------
Between EEG-R-Ant. and EEG-L-Post., p= 0.92656
N_EEG-R-Ant. = 8103, N_EEG-L-Post. = 6822
mean: EEG-R-Ant. = 5.8686,  SE: EEG-R-Ant. = 0.06276
mean: EEG-L-Post. = 5.8817,  SE: EEG-L-Post. = 0.066523
-------
Between EEG-R-Ant. and LFP, p= 3.8446e-07

N_EEG-R-Ant. = 8103, N_LFP = 6563
mean: EEG-R-Ant. = 5.8686,  SE: EEG-R-Ant. = 0.06276
mean: LFP = 5.473,  SE: LFP = 0.061067
-------
Between EEG-R-Post. and EEG-L-Post., p= 0.04757
N_EEG-R-Post. = 7264, N_EEG-L-Post. = 6822
mean: EEG-R-Post. = 6.235,  SE: EEG-R-Post. = 0.075723
mean: EEG-L-Post. = 5.8817,  SE: EEG-L-Post. = 0.066523
-------
Between EEG-R-Post. and LFP, p= 3.7584e-12
N_EEG-R-Post. = 7264, N_LFP = 6563
mean: EEG-R-Post. = 6.235,  SE: EEG-R-Post. = 0.075723
mean: LFP = 5.473,  SE: LFP = 0.061067
-------
Between EEG-L-Post. and LFP, p= 7.1787e-07
N_EEG-L-Post. = 6822, N_LFP = 6563
mean: EEG-L-Post. = 5.8817,  SE: EEG-L-Post. = 0.066523
mean: LFP = 5.473,  SE: LFP = 0.061067

**Test results for IS:**
Between EEG-L-Ant. and EEG-R-Ant., p= 1.3063e-66
N_EEG-L-Ant. = 10141, N_EEG-R-Ant. = 10294
mean: EEG-L-Ant. = 7.8359,  SE: EEG-L-Ant. = 0.078792
mean: EEG-R-Ant. = 6.2221,  SE: EEG-R-Ant. = 0.055506
-------
Between EEG-L-Ant. and EEG-R-Post., p= 5.7832e-70
N_EEG-L-Ant. = 10141, N_EEG-R-Post. = 9921
mean: EEG-L-Ant. = 7.8359,  SE: EEG-L-Ant. = 0.078792
mean: EEG-R-Post. = 6.2289,  SE: EEG-R-Post. = 0.057793
-------
Between EEG-L-Ant. and EEG-L-Post., p= 1.5251e-100
N_EEG-L-Ant. = 10141, N_EEG-L-Post. = 10243
mean: EEG-L-Ant. = 7.8359,  SE: EEG-L-Ant. = 0.078792
mean: EEG-L-Post. = 5.9007,  SE: EEG-L-Post. = 0.051058
-------
Between EEG-L-Ant. and LFP, p= 3.1836e-151
N_EEG-L-Ant. = 10141, N_LFP = 9807
mean: EEG-L-Ant. = 7.8359,  SE: EEG-L-Ant. = 0.078792
mean: LFP = 5.5457,  SE: LFP = 0.04734
-------
Between EEG-R-Ant. and EEG-R-Post., p= 0.48394
N_EEG-R-Ant. = 10294, N_EEG-R-Post. = 9921
mean: EEG-R-Ant. = 6.2221,  SE: EEG-R-Ant. = 0.055506
mean: EEG-R-Post. = 6.2289,  SE: EEG-R-Post. = 0.057793
-------
Between EEG-R-Ant. and EEG-L-Post., p= 3.1147e-05
N_EEG-R-Ant. = 10294, N_EEG-L-Post. = 10243
mean: EEG-R-Ant. = 6.2221,  SE: EEG-R-Ant. = 0.055506
mean: EEG-L-Post. = 5.9007,  SE: EEG-L-Post. = 0.051058
-------
Between EEG-R-Ant. and LFP, p= 1.621e-21
N_EEG-R-Ant. = 10294, N_LFP = 9807
mean: EEG-R-Ant. = 6.2221,  SE: EEG-R-Ant. = 0.055506
mean: LFP = 5.5457,  SE: LFP = 0.04734
-------
Between EEG-R-Post. and EEG-L-Post., p= 0.00065375

N_EEG-R-Post. = 9921, N_EEG-L-Post. = 10243
mean: EEG-R-Post. = 6.2289,  SE: EEG-R-Post. = 0.057793
mean: EEG-L-Post. = 5.9007,  SE: EEG-L-Post. = 0.051058
-------
Between EEG-R-Post. and LFP, p= 2.9154e-18
N_EEG-R-Post. = 9921, N_LFP = 9807
mean: EEG-R-Post. = 6.2289,  SE: EEG-R-Post. = 0.057793
mean: LFP = 5.5457,  SE: LFP = 0.04734
-------
Between EEG-L-Post. and LFP, p= 5.4635e-08
N_EEG-L-Post. = 10243, N_LFP = 9807
mean: EEG-L-Post. = 5.9007,  SE: EEG-L-Post. = 0.051058
mean: LFP = 5.5457,  SE: LFP = 0.04734

**2. Percentage**

**<u>Test results for SWS:</u>**
Between EEG-L-Ant. and EEG-R-Ant., p= 0.7
N_EEG-L-Ant. = 3, N_EEG-R-Ant. = 3
mean: EEG-L-Ant. = 22.7333,  SE: EEG-L-Ant. = 1.7362
mean: EEG-R-Ant. = 20.3667,  SE: EEG-R-Ant. = 2.9077
-------
Between EEG-L-Ant. and EEG-R-Post., p= 1
N_EEG-L-Ant. = 3, N_EEG-R-Post. = 3
mean: EEG-L-Ant. = 22.7333,  SE: EEG-L-Ant. = 1.7362
mean: EEG-R-Post. = 24.4333,  SE: EEG-R-Post. = 3.3933
-------
Between EEG-L-Ant. and EEG-L-Post., p= 0.2
N_EEG-L-Ant. = 3, N_EEG-L-Post. = 2
mean: EEG-L-Ant. = 22.7333,  SE: EEG-L-Ant. = 1.7362
mean: EEG-L-Post. = 32.4,  SE: EEG-L-Post. = 1.8
-------
Between EEG-L-Ant. and LFP, p= 0.1
N_EEG-L-Ant. = 3, N_LFP = 3
mean: EEG-L-Ant. = 22.7333,  SE: EEG-L-Ant. = 1.7362
mean: LFP = 32.0667,  SE: LFP = 1.593
-------
Between EEG-R-Ant. and EEG-R-Post., p= 0.4
N_EEG-R-Ant. = 3, N_EEG-R-Post. = 3
mean: EEG-R-Ant. = 20.3667,  SE: EEG-R-Ant. = 2.9077
mean: EEG-R-Post. = 24.4333,  SE: EEG-R-Post. = 3.3933
-------
Between EEG-R-Ant. and EEG-L-Post., p= 0.2
N_EEG-R-Ant. = 3, N_EEG-L-Post. = 2
mean: EEG-R-Ant. = 20.3667,  SE: EEG-R-Ant. = 2.9077
mean: EEG-L-Post. = 32.4,  SE: EEG-L-Post. = 1.8
-------
Between EEG-R-Ant. and LFP, p= 0.1
N_EEG-R-Ant. = 3, N_LFP = 3
mean: EEG-R-Ant. = 20.3667,  SE: EEG-R-Ant. = 2.9077
mean: LFP = 32.0667,  SE: LFP = 1.593
-------
Between EEG-R-Post. and EEG-L-Post., p= 0.4
N_EEG-R-Post. = 3, N_EEG-L-Post. = 2
mean: EEG-R-Post. = 24.4333,  SE: EEG-R-Post. = 3.3933
mean: EEG-L-Post. = 32.4,  SE: EEG-L-Post. = 1.8

-------
Between EEG-R-Post. and LFP, p= 0.4
N_EEG-R-Post. = 3, N_LFP = 3
mean: EEG-R-Post. = 24.4333,  SE: EEG-R-Post. = 3.3933
mean: LFP = 32.0667,  SE: LFP = 1.593
-------
Between EEG-L-Post. and LFP, p= 1
N_EEG-L-Post. = 2, N_LFP = 3
mean: EEG-L-Post. = 32.4,  SE: EEG-L-Post. = 1.8
mean: LFP = 32.0667,  SE: LFP = 1.593

**Test results for REM:**
Between EEG-L-Ant. and EEG-R-Ant., p= 0.4
N_EEG-L-Ant. = 3, N_EEG-R-Ant. = 3
mean: EEG-L-Ant. = 55,  SE: EEG-L-Ant. = 3.5119
mean: EEG-R-Ant. = 46,  SE: EEG-R-Ant. = 3.5119
-------
Between EEG-L-Ant. and EEG-R-Post., p= 0.1
N_EEG-L-Ant. = 3, N_EEG-R-Post. = 3
mean: EEG-L-Ant. = 55,  SE: EEG-L-Ant. = 3.5119
mean: EEG-R-Post. = 44.6667,  SE: EEG-R-Post. = 4.8419
-------
Between EEG-L-Ant. and EEG-L-Post., p= 0.8
N_EEG-L-Ant. = 3, N_EEG-L-Post. = 2
mean: EEG-L-Ant. = 55,  SE: EEG-L-Ant. = 3.5119
mean: EEG-L-Post. = 46,  SE: EEG-L-Post. = 8
-------
Between EEG-L-Ant. and LFP, p= 0.4
N_EEG-L-Ant. = 3, N_LFP = 3
mean: EEG-L-Ant. = 55,  SE: EEG-L-Ant. = 3.5119
mean: LFP = 40.3333,  SE: LFP = 6.3596
-------
Between EEG-R-Ant. and EEG-R-Post., p= 1
N_EEG-R-Ant. = 3, N_EEG-R-Post. = 3
mean: EEG-R-Ant. = 46,  SE: EEG-R-Ant. = 3.5119
mean: EEG-R-Post. = 44.6667,  SE: EEG-R-Post. = 4.8419
-------
Between EEG-R-Ant. and EEG-L-Post., p= 1
N_EEG-R-Ant. = 3, N_EEG-L-Post. = 2
mean: EEG-R-Ant. = 46,  SE: EEG-R-Ant. = 3.5119
mean: EEG-L-Post. = 46,  SE: EEG-L-Post. = 8
-------
Between EEG-R-Ant. and LFP, p= 0.5
N_EEG-R-Ant. = 3, N_LFP = 3
mean: EEG-R-Ant. = 46,  SE: EEG-R-Ant. = 3.5119
mean: LFP = 40.3333,  SE: LFP = 6.3596
-------
Between EEG-R-Post. and EEG-L-Post., p= 0.8
N_EEG-R-Post. = 3, N_EEG-L-Post. = 2
mean: EEG-R-Post. = 44.6667,  SE: EEG-R-Post. = 4.8419
mean: EEG-L-Post. = 46,  SE: EEG-L-Post. = 8
-------
Between EEG-R-Post. and LFP, p= 0.8
N_EEG-R-Post. = 3, N_LFP = 3
mean: EEG-R-Post. = 44.6667,  SE: EEG-R-Post. = 4.8419
mean: LFP = 40.3333,  SE: LFP = 6.3596

-------
Between EEG-L-Post. and LFP, p= 0.4
N_EEG-L-Post. = 2, N_LFP = 3
mean: EEG-L-Post. = 46,  SE: EEG-L-Post. = 8
mean: LFP = 40.3333,  SE: LFP = 6.3596

**Test results for IS:**
Between EEG-L-Ant. and EEG-R-Ant., p= 0.1
N_EEG-L-Ant. = 3, N_EEG-R-Ant. = 3
mean: EEG-L-Ant. = 22.3333,  SE: EEG-L-Ant. = 4.4096
mean: EEG-R-Ant. = 32.6667,  SE: EEG-R-Ant. = 2.6667
-------
Between EEG-L-Ant. and EEG-R-Post., p= 0.4
N_EEG-L-Ant. = 3, N_EEG-R-Post. = 3
mean: EEG-L-Ant. = 22.3333,  SE: EEG-L-Ant. = 4.4096
mean: EEG-R-Post. = 30.7,  SE: EEG-R-Post. = 7.6531
-------
Between EEG-L-Ant. and EEG-L-Post., p= 1
N_EEG-L-Ant. = 3, N_EEG-L-Post. = 2
mean: EEG-L-Ant. = 22.3333,  SE: EEG-L-Ant. = 4.4096
mean: EEG-L-Post. = 21.1,  SE: EEG-L-Post. = 9.9
-------
Between EEG-L-Ant. and LFP, p= 0.7
N_EEG-L-Ant. = 3, N_LFP = 3
mean: EEG-L-Ant. = 22.3333,  SE: EEG-L-Ant. = 4.4096
mean: LFP = 26.4,  SE: LFP = 5.5582
-------
Between EEG-R-Ant. and EEG-R-Post., p= 0.6
N_EEG-R-Ant. = 3, N_EEG-R-Post. = 3
mean: EEG-R-Ant. = 32.6667,  SE: EEG-R-Ant. = 2.6667
mean: EEG-R-Post. = 30.7,  SE: EEG-R-Post. = 7.6531
-------
Between EEG-R-Ant. and EEG-L-Post., p= 0.8
N_EEG-R-Ant. = 3, N_EEG-L-Post. = 2
mean: EEG-R-Ant. = 32.6667,  SE: EEG-R-Ant. = 2.6667
mean: EEG-L-Post. = 21.1,  SE: EEG-L-Post. = 9.9
-------
Between EEG-R-Ant. and LFP, p= 0.4
N_EEG-R-Ant. = 3, N_LFP = 3
mean: EEG-R-Ant. = 32.6667,  SE: EEG-R-Ant. = 2.6667
mean: LFP = 26.4,  SE: LFP = 5.5582
-------
Between EEG-R-Post. and EEG-L-Post., p= 0.8
N_EEG-R-Post. = 3, N_EEG-L-Post. = 2
mean: EEG-R-Post. = 30.7,  SE: EEG-R-Post. = 7.6531
mean: EEG-L-Post. = 21.1,  SE: EEG-L-Post. = 9.9
-------
Between EEG-R-Post. and LFP, p= 0.7
N_EEG-R-Post. = 3, N_LFP = 3
mean: EEG-R-Post. = 30.7,  SE: EEG-R-Post. = 7.6531
mean: LFP = 26.4,  SE: LFP = 5.5582
-------
Between EEG-L-Post. and LFP, p= 0.8
N_EEG-L-Post. = 2, N_LFP = 3
mean: EEG-L-Post. = 21.1,  SE: EEG-L-Post. = 9.9
mean: LFP = 26.4,  SE: LFP = 5.5582

**Congruence in each stage of sleep, averages over sites**

Congruence SWS: mean = 44.1667,  SD = 10.7266
-------
Congruence REM: mean = 54.5,  SD = 19.7967
-------
Congruence IS: mean = 47.8333,  SD = 9.3306
